# *commonPeak*: Equivalence testing to identify common ChIP-seq peaks across conditions and protocols

**DOI:** 10.64898/2026.02.16.706124

**Authors:** August H. Swillus, Francesca Tiso, Siddharth Annaldasula, Eldar Abdullaev, Regine Armann, Peter F. Arndt, Kirsten Kübler

## Abstract

**Background:** New ChIP-seq protocols are increasingly adopted alongside established workflows. However, dedicated methods are lacking to quantify the agreement of peak location and intensity across datasets, including comparisons across protocols and biological conditions.

**Objectives:** We present *commonPeak*, a statistical framework that identifies shared peaks across samples and tests whether their enrichment is similar across conditions, thereby supporting bench-marking of ChIP-seq protocols and cross-condition comparisons.

**Results:** *commonPeak* operates on BED peak sets and BAM files. As a use case, we applied it to an estrogen receptor alpha (ERα) ChIP-seq dataset from tamoxifen-sensitive and -resistant breast cancer cell line MCF-7 cells and identified 225 shared peaks with significantly similar enrichment. These peaks were largely distinct from differentially bound sites and enriched for estrogen signaling. This illustrates how equivalence-based peak selection can help separate conserved ERα-driven programs from condition-specific changes in ERα-targeting endocrine drugs such as tamoxifen.

**Availability and implementation:** *commonPeak* is freely available for non-commercial use via GitHub, with documentation and usage examples.

## 1 Introduction

Chromatin immunoprecipitation sequencing (ChIP-seq) is widely used to map transcription factor binding sites and histone modification enrichment (Kundaje *et al*. 2015). These epigenetic features are typically represented as genomic regions of significant ChIP-seq signal enrichment („peaks”). A large fraction of downstream analyses focuses either on identifying differences in binding/enrichment across conditions, such as DiffBind (Ross-Innes *et al*. 2012), csaw (Lun and Smyth 2016), and MAnorm (Shao *et al*. 2012), or on assessing replicate reproducibility within a condition, such as IDR (Jalili, Cremona and Palluzzi 2023).

Meanwhile, the methodological landscape is evolving rapidly. For example, modified versions of the ChIP-seq protocol are emerging to enable profiling from limited tissue material such as scarce *in vivo* hematopoietic cell populations or samples derived from small precancerous lesions (Lara-Astiaso *et al*. 2014). Benchmarking such approaches—and, more generally, comparing datasets across protocols or biological conditions— requires identifying peaks that agree in both genomic coordinates and signal strength, here referred to as “common peaks”, across experiments. In current practice, common peaks are often defined by coordinate overlap and the absence of a significant differential signal, but failure to detect a significant difference does not provide evidence that the same peak is consistently seen across experiments.

To address this gap, we introduce *commonPeak*, an agreement-testing method that operationalizes common peaks through a two-stage definition: (i) shared presence across samples and (ii) statistical equivalence of peak strength. This framework enables positive conclusions of concordant enrichment across conditions and protocols, making it particularly suited for benchmarking new ChIP-seq protocols against established standards and for identifying conserved binding programs across conditions.

## 2 Methods

### 2.1 Inputs

To quantify and test concordance of ChIP-seq peak intensities (Fig. 1), *commonPeak* takes (i) a set of peak intervals in BED format for each sample and (ii) aligned reads in BAM format from all tested samples, including input control samples.

**Figure 1.**
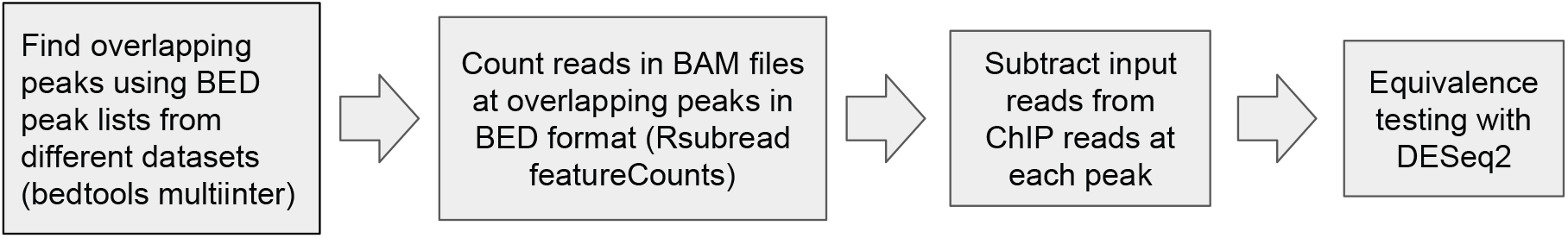
Schematic overview of the *commonPeak* workflow.

### 2.2 Defining and read counting in intervals

Because *commonPeak* is designed for benchmarking and cross-condition concordance testing, candidate regions are restricted to peaks that are present in all samples. The first step in the pipeline thus identifies the intersection of peak ranges using bedtools *multiinter* (Quinlan and Hall 2010). Only intervals with support in all samples are retained as candidates for subsequent intensity testing. *commonPeak* calculates the average coverage signal in the peak intervals (default: 400 bp wide in narrow mode; can be set to 1000 bp in broad mode) centered around the point of the highest read pile-up in each peak that overlaps between all samples from all conditions.

Next, the reads overlapping the candidate intervals are counted for each sample using Rsubread’s *featureCounts* function (Liao, Smyth and Shi 2014). This produces a matrix in which each row is a peak and each column represents a sample, with values representing the number of read counts falling into this peak.

### 2.3 Statistical testing of peak strength

To test peak strength, *commonPeak* leverages DESeq2, which expects unnormalized integer count matrices as input (Love, Huber and Anders 2014). ChIP-seq experiments typically include input control samples to account for unspecific antibody binding and artifacts associated with sample preparation (Kundaje *et al*. 2015). Thus, prior to DESeq2, *commonPeak* subtracts input read counts from ChIP counts in each peak, analogous to the DiffBind workflow, which also provides read counts in ChIP peaks to DESeq2, albeit exclusively for differential binding analysis (Ross-Innes *et al*. 2012). Before subtraction, the input read counts are scaled by the ratio of the library size of each ChIP sample to the library size of its respective input control sample to account for differences in sequencing depth across samples. Scaled values are then rounded to the nearest integer. All negative values after subtraction are set to zero.

The final test of whether peak enrichment is equivalent between conditions is performed using the argument *altHypothesis = “lessAbs”* from DESeq2, which fits a negative binomial model to the read counts from all samples, including all replicates, and estimates a log_2_ fold change (log_2_FC) between conditions for each peak. Equivalence is then assessed using a two one-sided Wald test (TOST) procedure (Schuirmann 1987; Love, Huber and Anders 2014). Contrary to traditional hypothesis tests, these TOSTs test two null hypotheses for each peak (i.e., asking whether the log_2_FC is sufficiently close to zero to be considered similar). Specifically, the tests determine whether the absolute value of the log_2_FC is smaller than a threshold for functional equivalence of peak strength (Δ) set by the user (Schuirmann 1987):

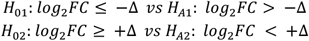

The two one-sided tests calculate the test statistics *t*_*1*_ and *t*_*2*_, taking the standard error (SE) into account:

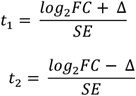

Only if both null hypotheses are rejected is a peak regarded as significantly similar between conditions within the tolerance of Δ. Multiple testing is controlled by DESeq2 using the Benjamini–Hochberg procedure.

### 2.4 Outputs and visualization

*commonPeak* outputs (i) a summary table of the equivalence testing result, (ii) a file of common peak intervals, and (iii) MA plots generated using the *plotMA*() function from DESeq2 to visualize the log_2_FC between conditions against the average read count for all peaks with significantly similar intensity across conditions.

### 2.5 Auxiliary analyses used in this study

For benchmarking *commonPeak*, we used estrogen receptor alpha (ERα) ChIP-seq data from the ERα-positive breast adenocarcinoma cell line MCF-7 (Ross-Innes *et al*. 2012), including three tamoxifen-responsive replicates (GSM798423, GSM798424, GSM798425), two tamoxifen-resistant replicates (GSM798432, GSM798433), and input controls (GSM798440, GSM798444).

Using the coverage matrix as input, we applied the *plotProfile* function from *deepTools* (Ramírez *et al*. 2016) with default settings to visualize signal profiles over *commonPeak* intervals. To account for differences in sequencing depth across samples, we used BPM (bins per million mapped reads)-normalized BigWig files, where the number of reads in each bin is divided by the total number of reads per sample. In this analysis, we relied on narrow 20-bp-wide bins to ensure accurate normalization across genomic regions with varying coverage. The intersection analysis between significantly similar peaks and significantly differential peaks identified by DiffBind was performed using BEDTools *intersect*. We used *Enrichr* to perform pathway enrichment analysis using the KEGG 2019 human gene set library (Chen *et al*. 2013).

## 3 Results

We benchmarked *commonPeak* on a well-characterized ERα ChIP-seq dataset from the breast cancer cell line MCF-7 comparing tamoxifen-responsive and tamoxifen-resistant conditions. Tamoxifen is an ERα-targeting endocrine drug that is widely used in ER-positive breast cancer. This setting provides a realistic test case in which ER binding is expected to be conserved across conditions, alongside a set of condition-specific changes, enabling evaluation of *commonPeak*’s ability to identify peaks that are shared in genomic location and comparable in strength across conditions. *commonPeak* runs in 3 minutes on 5 samples and 27,601 peaks, using BAM files from 715 MB to 999 MB in size. Peaks were classified as similar if |log_2_FC| < 0.75 with a *q* < 0.05. In contrast, peaks with a |log_2_ FC| > 0.75 and a *q* < 0.05 were defined as differential within the *commonPeak* analysis. To detect peaks with significantly decreased or increased intensities in tamoxifen-resistant versus tamoxifen-sensitive samples, we used DESeq2, specifying altHypothesis = “greater” or “less”, respectively.

At our significance threshold, we identified 225 peaks with similar ERα binding and 4,546 peaks with differential binding out of 27,601 tested peaks. The set of peaks called equivalent by *commonPeak* between the resistant and responsive conditions did not overlap with peaks called differential when using a separate differential binding analysis with DiffBind (*q* = 0.05, Benjamini–Hochberg-corrected), consistent with the opposing definitions of equivalence and differential binding. Compared with the ChIP-seq signal across all peak intervals identified in all samples (Fig. 2A), peaks classified as statistiocally equivalent showed log_2_FC around zero and higher average intensities (BPM-normalized mean 5.6 ± 0.86; Fig. 2B) than peaks upregulated in tamoxifen-resistant samples (BPM-normalized mean 3.28 ± 0.92; Fig. 2C) and peaks downregulated in ta-moxifen-resistant samples (BPM-normalized mean 3.83 ± 1.09; Fig. 2D), suggesting that *commonPeak* prefentially identifies a high-confidence subset of strong ERα-binding sites whose enrichment is maintained across conditions.

**Figure 2.**
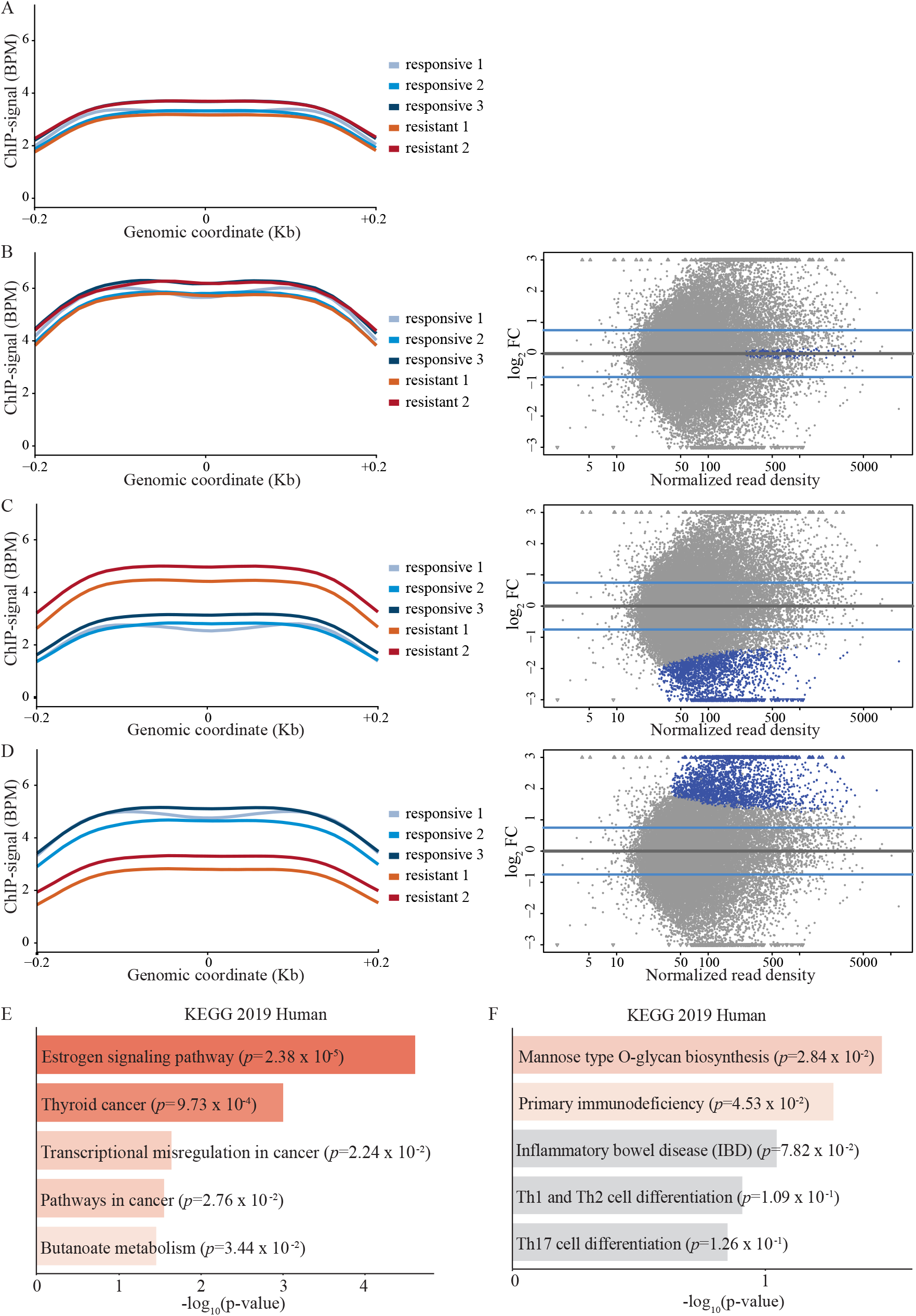
Significantly similar peaks are associated with log_2_FC centered around zero and high signal intensity. **A-D, left**. deepTools plotProfile plots of the average BPM-normalized ChIP-seq signal across (**A**) all peak intervals present in all samples, (**B**) all significantly similar peaks identified by *commonPeak*, (**C**) peaks classified as differential in resistant samples, and (**D**) peaks classified as differential in responsive samples. The x-axis is centered on the peak midpoint (zero) of all peaks; each line represents the BPM-normalized ChIP-seq signal for one sample (color-coded). **A-D, right**. MA plots for the same peak sets. The y-axis shows the log_2_FC between conditions (tamoxifen-resistant vs. tamoxifen-responsive), the y-axis the mean normalized signal. In **B** significantly similar peaks are marked in blue with log_2_FC around zero and high read counts; blue dots indicate differental peaks with low read counts in resistant (**C**) and responsive (**D**) conditions, respectively. **E, F**. Pathway enrichment for genes assigned to peaks. Genes were defined as proximal if their TSS lay within ±20 kb of the peak center. Enrichment was computed using Enrichr (KEGG 2019 Human) for (**E**) 25 genes assigned to the most significant equivalent peaks and (**F**) 25 genes assigned to differential peaks. Bar plots show enrichment significance (Fisher’s exact test *p*-values); red indicates significant terms (darker shades indicate smaller *p*-values) and grey indicates non-significant terms.

We therefore hypothesized that these sites would correspond to a core ERα regulatory program independet of tamoxifen response and would occur preferentially near genes associated with canonical estrogen signaling. To test this hypothesis, we associated common and differential peaks with nearby genes by assigning transcription start sites (TSS) within ±20 kb of each peak center within the 400-bp-wide peaks. Next, we performed pathway enrichment analyses using 25 unique genes assigned to common peaks and, to match gene set sizes, 25 genes randomly sampled from tjose assigned to differential peaks. Genes proximal to the common peaks showed significant enrichment for the KEGG 2019 estrogen signaling pathway gene set (*p* = 5.72 x 10^-4^, Benjamini–Hochberg-corrected Fisher’s exact test; Fig. 2E, Table 1), consistent with a core ERα regulatory program shared across conditions. In contrast, genes proximal to differential peaks identified by DiffBind were not enriched for the estrogen pathway and showed only weak and heterogenous enrichments (Fig. 2F, Table 1). Together, these finding support the biological interpretation that *common-Peak*-equivalent ERα sites are enriched near canonical estrogen signaling genes, consistent with a core ERα-binding program maintained across tamoxifen-sensitive and -resistant states, whereas condition-specific ERα-binding changes are more dispersed and show weaker, heterogeneous pathway enrichment.

**Table 1.** Pathway enrichment analysis of genes proximal to common peaks and differential peaks. Enrichr results (KEGG 2019 Human) for 25 genes assigned to common peaks (top) and differential peaks (bottom). Shown are *p*-value from Fisher’s exact test, Benjamini–Hochberg-corrected *p*-value, odds ratio, and Enrichr combined score for the most strongly enriched gene sets (adj, adjusted; OR, odds ratio; comb, combined)

| Pathway enrichment for 25 genes proximal to common peaks |  |  |  |  |
| --- | --- | --- | --- | --- |
| Term | *p*-value | Adj. *p*-value | OR | Comb. Score |
| Estrogen signaling pathway | $2.38 \times 10^{-5}$ | $5.72 \times 10^{-4}$ | 28.417 | 302.479 |
| Thyroid cancer | $9.73 \times 10^{-4}$ | 0.012 | 49.540 | 343.590 |
| Transcriptional misregulation in cancer | 0.022 | 0.125 | 9.353 | 35.517 |
| Pathways in cancer | 0.028 | 0.125 | 5.032 | 18.064 |
| Butanoate metabolism | 0.034 | 0.125 | 30.784 | 103.698 |

**Table 1. Pathway enrichment analysis of genes proximal to common peaks and differential peaks.** Enrichr results (KEGG 2019 Human) for 25 genes assigned to common peaks
| Pathway enrichment of 25 genes proximal to differential peaks |  |  |  |  |
| --- | --- | --- | --- | --- |
| Term | *p*-value | Adj. *p*-value | OR | Comb. Score |
| Mannose type O-glycan biosynthesis | $2.84 \times 10^{-2}$ | 0.317 | 37.79 | 134.619 |
| Primary immunodeficiency | 0.045 | 0.317 | 23.078 | 71.431 |
| Inflammatory bowel disease | 0.078 | 0.318 | 12.963 | 33.035 |
| Th1 and Th2 cell differentiation | 0.109 | 0.3181 | 9.104 | 20.185 |
| Th17 cell differentiation | 0.126 | 0.318 | 7.81 | 16.205 |

## 4 Conclusion

The here presented *commonPeak* provides a user-friendly workflow for identifying genomic regions that are shared across samples and statistically similar in binding strength across conditions. Users only need to run a single command to obtain a list of common peak regions with associated *q-*values, as well as MA plots for visualization. In an ERα ChIP-seq bench-marking analysis, *commonPeak* recovered a set of equivalently bound sites that was distinct from differentially bound peaks and was enriched near genes involved in estrogen signaling, supporting its use for cross-condition concordance analyses.

*commonPeak* currently prioritizes high-confidence agreement by requiring peaks to be present in all samples, a conservative design choice to ensure comparability between experiments. Future versions could offer users to relax this requirement and test for significant similarity at genomic locations that failed peak calling in one sample due to calling variability. Additional limitations include restricted normalization options (e.g., no spike-in normalization) and reliance on DESeq2 for equivalence testing. In future extensions, *commonPeak* could be also further adapted to be compatible with chromatin accessibility data, such as from ATAC-seq experiments, additional count-based testing backends beyond DESeq2 (e.g., edgeR), and richer design matrices.

Together, *commonPeak* offers a reproducible equivalence-based route to identifying and reporting concordant peaks for downstream biological interpretation or ChIP-seq method comparison.

## Author contributions

A.H.S.: Conceptualization, Methodology, Software, Formal analysis, Visualization, Writing—original draft. F.T.: Conceptualization, Methodology, Formal analysis, Writing— original draft. E.A.: Software, Writing—review & editing. S.A.: Software, Writing—review & editing. R.A.: Writing—review & editing. P.F.A.: Software, Writing—review & editing. K.K.: Conceptualization, Methodology, Supervision, Funding acquisition, Writing—review & editing.

## Code and data availability

The *commonPeak* package will be made publicly accessible on GitHub. The benchmarking data are available from the Gene Express Omnibus (GEO) under the accession number GSE32222.

